# Molecular analysis of a public cross-neutralizing antibody response to SARS-CoV-2

**DOI:** 10.1101/2022.05.17.492220

**Authors:** Meng Yuan, Yiquan Wang, Huibin Lv, Ian A. Wilson, Nicholas C. Wu

## Abstract

As SARS-CoV-2 variants of concerns (VOCs) continue to emerge, cross-neutralizing antibody responses become key towards next-generation design of a more universal COVID-19 vaccine. By analyzing published data from the literature, we report here that the combination of germline genes IGHV2-5/IGLV2-14 represents a public antibody response to the receptor-binding domain (RBD) that potently cross-neutralizes all VOCs to date, including Omicron and its sub-lineages. Detailed molecular analysis shows that the complementarity-determining region H3 sequences of IGHV2-5/IGLV2-14-encoded RBD antibodies have a preferred length of 11 amino acids and a conserved HxIxxI motif. In addition, these antibodies have a strong allelic preference due to an allelic polymorphism at amino-acid residue 54 of IGHV2-5, which locates at the paratope. These findings have important implications for understanding cross-neutralizing antibody responses to SARS-CoV-2 and its heterogenicity at the population level as well as the development of a universal COVID-19 vaccine.

## MAIN

The effectiveness of COVID-19 vaccines has been challenged by the evolution of diverse SARS-CoV-2 variants in the past two years. The recent emergence of Omicron and its sub-lineages BA.2, BA.2.12.1, BA.4, and BA.5 further highlights the urgent need for a more broadly protective vaccine. An ideal COVID-19 vaccine should elicit high titers of neutralizing antibodies that are potent against antigenically distinct variants. However, many potent neutralizing antibodies only have limited cross-reactivity for variants other than the immunizing strain. For example, a major class of antibodies to the receptor-binding domain (RBD) that are encoded by IGHV3-53/3-66 are highly potent against the ancestral Hu-1 strain, but most of them lose their activity against many other variants [1, 2]. Similarly, Beta-specific antibodies can be elicited without cross-neutralizing activity against ancestral or other variants [3]. On the other hand, antibodies to S2 are typically broadly reactive but have weak neutralizing activity [4-6]. Nevertheless, a few RBD antibodies exhibit marked neutralization potency and breadth, as exemplified by those to the RBS-D epitope [2].

One representative RBS-D antibody is LY-CoV1404 (also known as Bebtelovimab), which is a monoclonal therapeutic antibody from Eli Lilly. LY-CoV1404 is encoded by IGHV2-5/IGLV2-14 and can cross-neutralize the ancestral Hu-1 strain as well as all known variants of concern (VOCs), including Omicron and circulating sub-lineages [7, 8]. In fact, the binding mode of LY-CoV1404 is identical to the cross-neutralizing antibody 2-7, which is also encoded by IGHV2-5/IGLV2-14 [9]. More recently, Veesler and colleagues reported another potently cross-neutralizing antibody with similar sequences and binding mode as LY-CoV1404 [10]. As IGHV2-5 was shown to be an important contributor to cross-neutralizing antibody response [11], the observations above stimulated a systematic analysis of IGHV2-5/IGLV2-14-encoded RBD antibodies to SARS-CoV-2.

In our previous study, we assembled a dataset of ∼8,000 antibodies to SARS-CoV-2 spike (S) protein [12]. This dataset contains seven IGHV2-5/IGLV2-14-encoded RBD antibodies, including LY-CoV1404, from six different donors [7, 13-17]. In addition, four additional IGHV2-5/IGLV2-14-encoded RBD antibodies were reported in a recent study [18]. Our analysis here is therefore based on a total of 11 IGHV2-5/IGLV2-14-encoded RBD antibodies from at least seven donors. Three of these 11 antibodies have available information for the complete nucleotide sequence, nine have complete amino-acid sequence information, 10 have amino-acid sequence information for the complementarity-determining regions (CDRs) H3 and L3, and four have structure information. Neutralizing data from previous studies have demonstrated that these IGHV2-5/IGLV2-14-encoded RBD antibodies have high cross-neutralizing activity [7, 18-20], some of which remain potent against Omicron (**Figure 1A**). Previous studies have also shown that they compete with ACE2 for RBD binding [7, 18, 20, 21] (**Figure S1**).

**Figure 1.**
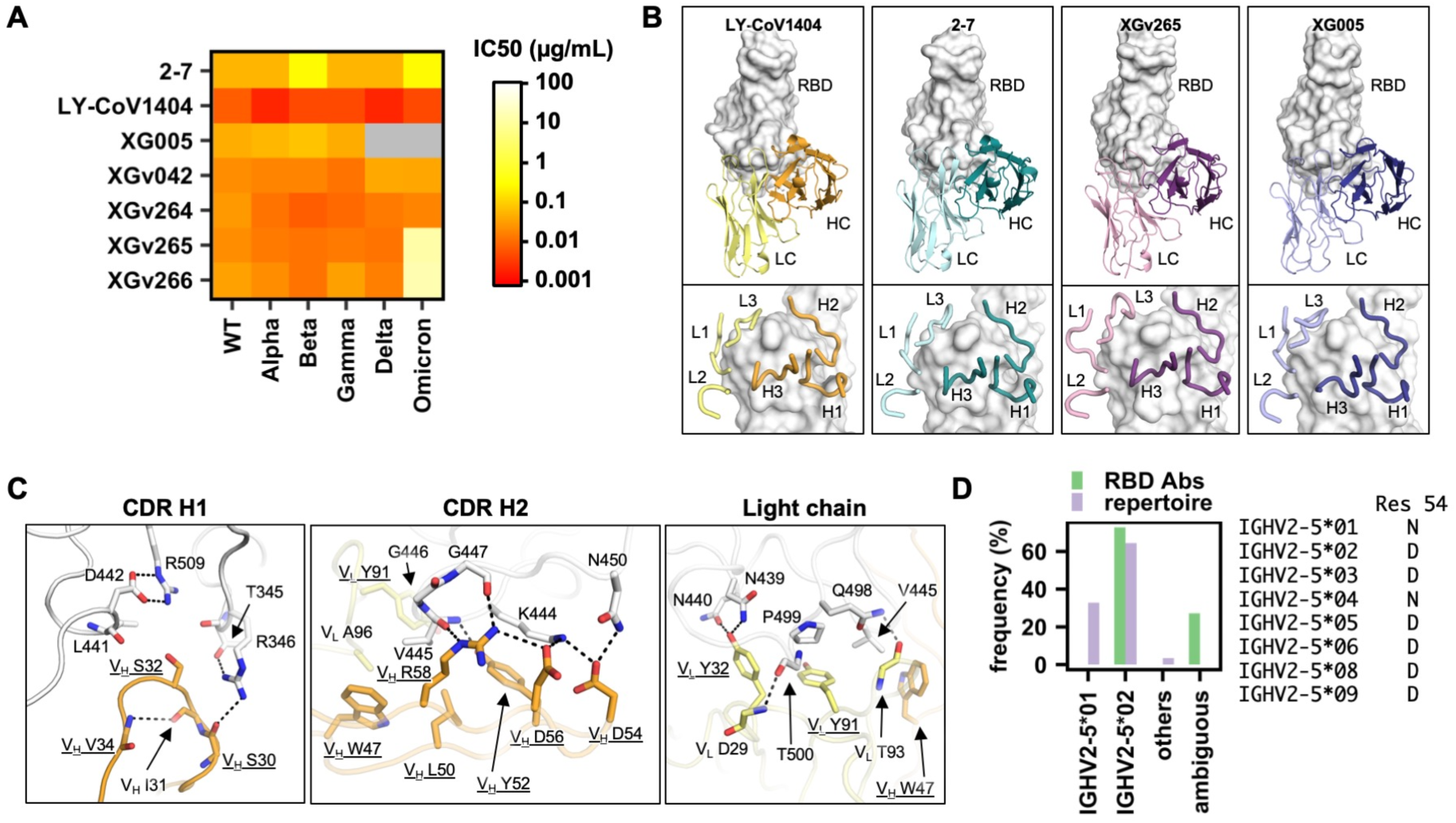
IGHV2-5/IGLV2-14 is a public antibody response with strong allelic preference. **(A)** The half maximal inhibitory concentration (IC_50_) of IGHV2-5/IGLV2-14-encoded RBD antibodies against different SARS-CoV-2 VOCs in pseudovirus assays. Data were taken from previous studies [7, 18-20]. **(B)** Four IGHV2-5/IGLV2-14-encoded RBD antibodies have structure information available. Their binding modes to RBD (white surface) are compared. Upper panels: heavy chain (HC) and light chain (LC) of each antibody are shown. Bottom panels: zoom-in views with six CDR loops of each antibody shown. LY-CoV1404: PDB 7MMO [7]. 2-7: PDB 7LSS [21]. XGv265: PDB 7WEE [18]. XG005: PDB 7V26 [20]. **(C)** Key interactions between LY-CoV1404 and RBD are shown. Hydrogen bonds and salt bridges are represented by black dashed lines. All germline-encoded residues are underlined. Heavy chain is in orange, light chain in yellow, and RBD is in white. **(D)** IGHV allele usage of the 11 IGHV2-5/IGLV2-14-encoded RBD antibodies (RBD Abs) is compared with that of IGHV2-5-encoded antibodies in published repertoire sequencing datasets from 13 healthy donors [22, 23]. The amino-acid identity at residue 54 of each IGHV2-5 allele is indicated.

Next, we performed a structural analysis to uncover the sequence determinants of IGHV2-5/IGLV2-14-encoded antibodies for RBD engagement. For antibody residues, the Kabat numbering scheme is used unless otherwise stated. All four IGHV2-5/IGLV2-14-encoded RBD antibodies with available structural information exhibit the same binding mode to the RBD (**Figure 1B**). As observed in LY-CoV1404, most amino-acid side chains in the paratope are germline-encoded and form key interactions with the RBD (**Figure 1C**). For example, germline-encoded V_H_ S32 in the CDR H1 fits into a polar pocket in the RBD. In addition, germline-encoded V_H_ Y52, D54, D56, and R58 in the CDR H2 form an extensive network of H-bonds and electrostatic interactions with the RBD. Furthermore, two key paratope residues in the light chain V_L_ Y32 and Y91 are also germline-encoded. These observations demonstrate that the RBD-binding determinants are encoded in the germline sequences of IGHV2-5 and IGLV2-14. Consistently, several IGHV2-5/IGLV2-14-encoded RBD antibodies have very few somatic hypermutations (SHMs) (**Table S1**).

For example, S24-223 has only one SHM, and COV2-2268 and 2-7 have only four each. Of note, none of their SHMs overlap.

Additional sequence analysis indicated that IGHV2-5/IGLV2-14-encoded RBD antibodies had a strong allelic preference towards IGHV2-5*02. Eight out of 11 IGHV2-5/IGLV2-14-encoded RBD antibodies could be assigned to IGHV2-5*02, while the allele usage for the other three was ambiguous (**Figure 1D and Table S1**). In contrast, analysis of the B cell repertoire in 13 healthy donors [22, 23] showed that alleles IGHV2-5*01 and IGHV2-5*02 were both commonly used, with a frequency of 33% and 64%, respectively, among all IGHV2-5 antibodies (**Figure 1D and Table S2**). The lack of IGHV2-5*01 among IGHV2-5/IGLV2-14-encoded RBD antibodies is likely due to an allelic polymorphism at residue 54. IGHV2-5*01 and IGHV2-5*02 have Asn and Asp, respectively, at residue 54. V_H_ D54 in IGHV2-5/IGLV2-14-encoded RBD antibodies plays an important role in RBD binding through a salt bridge with RBD K444 and an H-bond with RBD N450 (**Figure 1C**). Replacing the Asp at V_H_ residue 54 by Asn would convert the salt bridge with RBD K444 to a H-bond, which would likely significantly weaken the binding energy. Consistently, all eight of the nine IGHV2-5/IGLV2-14-encoded RBD antibodies with sequence information available have an Asp at V_H_ residue 54, whereas the remaining one has a Glu at V_H_ residue 54 (**Table S1**). These findings provide a mechanistic basis for the allelic preference against IGHV2-5*01, despite its prevalence in the human population. Coincidentally, an almost identical observation was observed in an IGHV2-5-encoded HIV antibody, in which V_H_ D54 results in much stronger binding than V_H_ N54 [24].

Lastly, we analyzed the CDR H3 sequences of the IGHV2-5/IGLV2-14-encoded RBD antibodies. Among 10 IGHV2-5/IGLV2-14-encoded RBD antibodies with CDR H3 sequence information available, eight had a CDR H3 length of 11 amino acids (IMGT numbering) and came from at least five patients (**Figure 2A**). The CDR H3 sequences from these eight antibodies shared a motif HxIxxI or conserved variations of it, including HxIxxL and HxVxxI (**Figures 2A and 2B**). The HxIxxI motif consisted of V_H_ H95, I97, and I100 (Kabat numbering) and is uncommon among the CDR H3 sequences of IGHV2-5-encoded antibodies in the human antibody repertoire (**Figure 2B**). V_H_ H95, I97, and I100 in the HxIxxI motif play critical roles in stabilizing the loop conformation as well as RBD binding (**Figure 2C**). V_H_ H95 forms two intramolecular H-bonds to stabilize the CDR H3 loop. The first H-bond involves the side chain of V_H_ Y52, which in turn H-bonds with RBD V445 amide nitrogen. The second H-bond involves the backbone carbonyl of V_H_ I100. In addition, V_H_ H95 also forms van der Waals interaction with RBD V445. V_H_ I97 at the tip of the CDR H3 loop inserts into a hydrophobic pocket formed by RBD V445 and P499, as well as the aliphatic portion of RBD N440. V_H_ I100 helps position V_L_ Y91 to interact with RBD V445 and P499. As shown by IgBlast analysis [25], the HxIxxI motif is largely encoded by N-nucleotide addition, although V_H_ I97 may sometimes be encoded by an IGHD gene (**Figure 2D**). Of note, while CDR H3 of XG005 has 12 amino acids (**Figure 2A**), it adopts a similar conformation to those with 11 amino acids (**Figure S2**). Overall, IGHV2-5/IGLV2-14-encoded RBD antibodies with a CDR H3 length of 11 amino acids have convergent CDR H3 sequences, and thus can be classified as a public clonotype.

**Figure 2.**
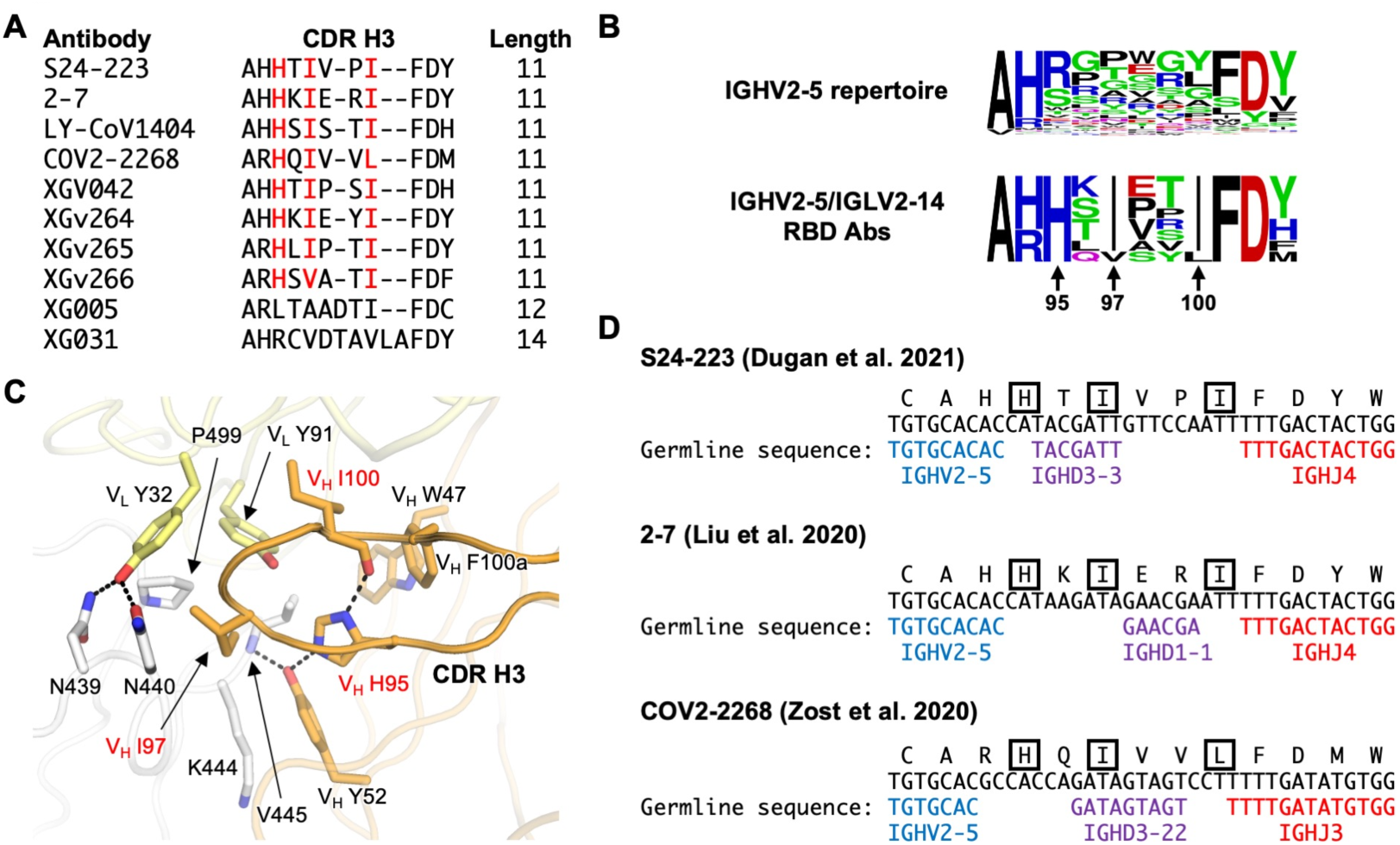
HxIxxI is a common motif in IGHV2-5/IGLV2-14-encoded RBD antibodies. **(A)** CDR H3 sequences (IMGT numbering) from IGHV2-5/IGLV2-14-encoded RBD antibodies are aligned. Residues of interest are highlighted in red. **(B)** CDR H3 sequences (IMGT numbering) of IGHV2-5/IGLV2-14-encoded RBD antibodies and IGHV2-5-encoded antibodies in the human antibody repertoire are shown as sequence logos. Only those antibodies with a CDR H3 length of 11 amino acids are included in this analysis. Residues of interest are labeled. Sequences of IGHV2-5-encoded antibodies in the human antibody repertoire were downloaded from the Observed Antibody Space [41]. A total of 9,197 IGHV2-5-encoded antibodies in the human antibody repertoire were analyzed here. Of note, while Kabat numbering was used for the residue position, IMGT numbering was used for defining CDR H3 length. **(C)** Interaction between the CDR H3 of LY-CoV1404 and RBD is shown. PDB 7MMO [7]. Hydrogen bonds are represented by black dashed lines. Heavy chain is in orange, light chain in yellow, and RBD is in white. Residues of interest are highlighted in red. **(D)** Amino-acid and nucleotide sequences of the V-D-J junction of three IGHV2-5/IGLV2-14-encoded RBD antibodies are shown. Putative germline sequences and segments are indicated. Residues of interest are boxed.

Due to the continuous evolution of SARS-CoV-2 VOCs, identification of cross-neutralizing human monoclonal antibodies has been a global research focus. IGHV1-58/IGKV3-20-encoded RBD antibodies are perhaps the most well-characterized public antibody clonotype that is cross-neutralizing against multiple SARS-CoV-2 VOCs [3, 12, 26-29]. However, recent studies have shown that many IGHV1-58/IGKV3-20-encoded RBD antibodies have minimal neutralizing activity against Omicron and its sub-lineages due to mutations Q493R and F486V on the RBD [30-32]. In comparison, IGHV2-5/IGLV2-14-encoded RBD antibodies, which mostly retain potency against Omicron and its sub-lineages (**Figure 1A**) [7, 8, 18-20, 33], have higher neutralization breadth. Since IGHV2-5/IGLV2-14-encoded RBD antibodies are also a public antibody clonotype, they further substantiate the rationale and strategy for development of a more universal COVID-19 vaccine.

Nevertheless, some individuals may have difficulties generating an IGHV2-5/IGLV2-14-encoded RBD antibody response, due to the alleles that they possess (**Figure 1D**). Since there is no known copy number variation for IGHV2-5 [34], each person should carry two copies of IGHV2-5 in the genome. If both copies are IGHV2-5*01 allele, the person may not have the suitable B cell germline clone to produce a IGHV2-5/IGLV2-14-encoded RBD antibody response. In fact, donor 112 in the 13 healthy donors that were analyzed in this study is very likely to be IGHV2-5*01 homozygous, since 94% of its IGHV2-5-encoded antibodies were assigned to IGHV2-5*01 (**Table S2**). Moreover, the conserved HxIxxI motif in CDR H3 of IGHV2-5/IGLV2-14-encoded RBD antibodies is mostly encoded by random N-nucleotide addition. As a result, B cell germline clones that can produce IGHV2-5/IGLV2-14-encoded RBD antibodies may be relatively rare. These results may provide a genetic basis for heterogenicity in the cross-neutralizing antibody response among different individuals. While allelic preference has previously been described for neutralizing antibodies to other viruses [24, 35-37], its clinical implications for COVID-19 remain to be fully explored.

## MATERIALS AND METHODS

### Dataset collection

The information on antibodies S24-223, P2B-1E4, 2-7, LY-CoV1404, XG005, XG031, and COV2-2268 were compiled in our previous study [12], whereas the information on XGv042, XGv264, XGv265, and XGv266 were compiled in CoV-AbDab [38]. Neutralization data of each monoclonal antibody were collected from the original papers (**Table S1**). Somatic hypermutations were identified by IgBlast [25].

### Allele assignment of IGHV2-5/IGLV2-14-encoded RBD antibodies

For antibodies P2B-1E4, XG005, and XG031, the allele information was obtained from the original publications [14, 16]. For other antibodies, IgBlast was used to assign the allele of each antibody [25]. Nucleotide sequence, if available, was used as input for IgBlast. Otherwise, protein sequence was used. If an antibody showed equally likely to be encoded by two or more alleles, the allele assignment would be classified as “ambiguous”. All “ambiguous” allele assignments in this study came from antibodies that do not have nucleotide sequence information available, namely XGv264, XGv265, and XGv266. Of note, while IgBlast showed that XGv266 was equally likely to be encoded by IGHV2-5*01 and IGHV2-5*02, we postulated that XGv266 should be assigned to IGHV2-5*02 at the nucleotide level. Specifically, XGv266 had a Glu at V_H_ residue 54, which was one nucleotide change from the Asp codon used (IGHV2-5*02) but two nucleotides away from Asn (IGHV2-5*01). However, IgBlast did not utilize codon information for allele assignment when the amino-acid sequence was used as input.

### Analysis of allele usage in published antibody repertoire

Published antibody repertoire sequencing datasets from 13 healthy donors [22, 23] were downloaded from cAb-Rep [39]. Putative germline gene alleles for each antibody sequence in these repertoire sequencing datasets from healthy donors were identified by IgBLAST [25].

### Analysis of CDR H3 sequences

Sequence alignment was performed using MAFFT [40]. Antibody sequences in the human antibody repertoire were downloaded from the Observed Antibody Space [41]. IGHV2-5 antibodies as well as their CDR H3 sequences were identified using IgBLAST [25]. Sequence logos were generated by WebLogo [42]. Putative germline sequences and segments in the V-D-J junctions were identified by IgBLAST [25].

### Code Availability

Custom codes for all analyses have been deposited to https://github.com/nicwulab/IGHV2-5_RBD_Abs.

## ACKNOWLEDGEMENTS

This work was supported by National Institutes of Health (NIH) DP2 AT011966 (N.C.W.), R01 AI167910 (N.C.W.), the Michelson Prizes for Human Immunology and Vaccine Research (N.C.W.), Bill and Melinda Gates Foundation INV-004923 (I.A.W.), and a Calmette and Yersin scholarship from the Pasteur International Network Association (H.L.).

## AUTHOR CONTRIBUTIONS

M.Y. and N.C.W. conceived and designed the study. All authors performed data analysis. M.Y. and N.C.W. wrote the paper and all authors reviewed and/or edited the paper.

---

**Figure S1.**
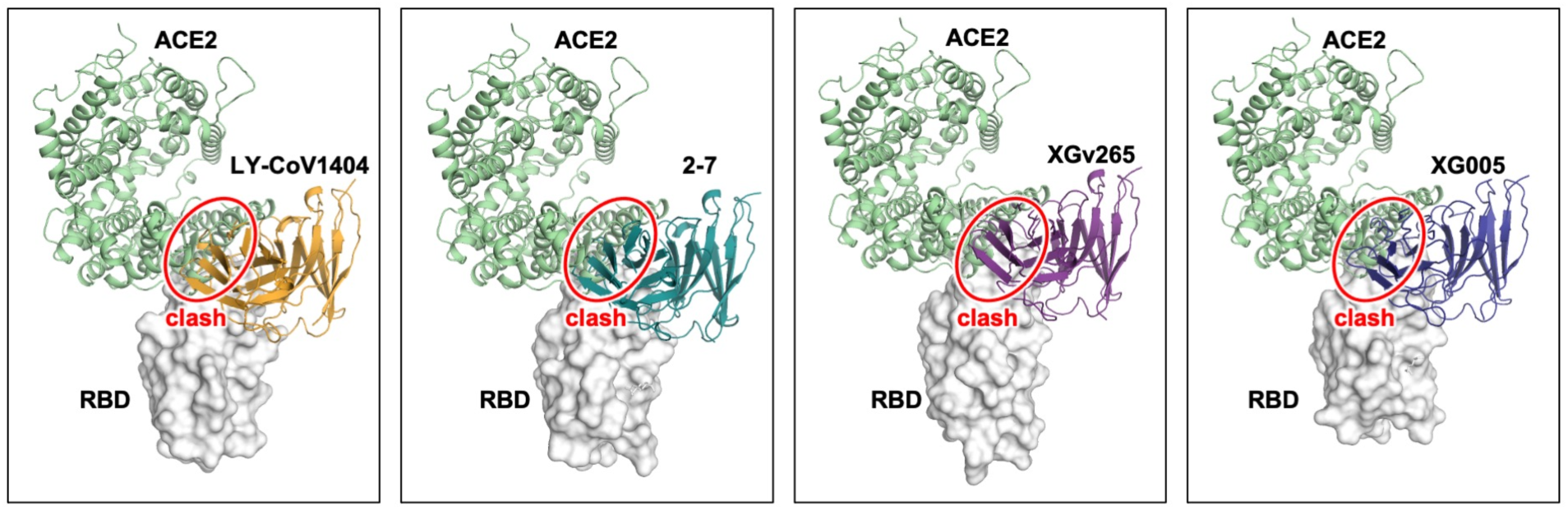
IGHV2-5/IGLV2-14 antibodies would clash with ACE2 binding. Structures of antibody/RBD complexes are superimposed onto the RBD/ACE2 complex structure (PDB 6M0J) [43]. RBD is represented by a white surface. ACE2 is shown as green cartoon. LY-CoV1404: PDB 7MMO [7]. 2-7: PDB 7LSS [21]. XGv265: PDB 7WEE [18]. XG005: PDB 7V26 [20].

**Figure S2.**
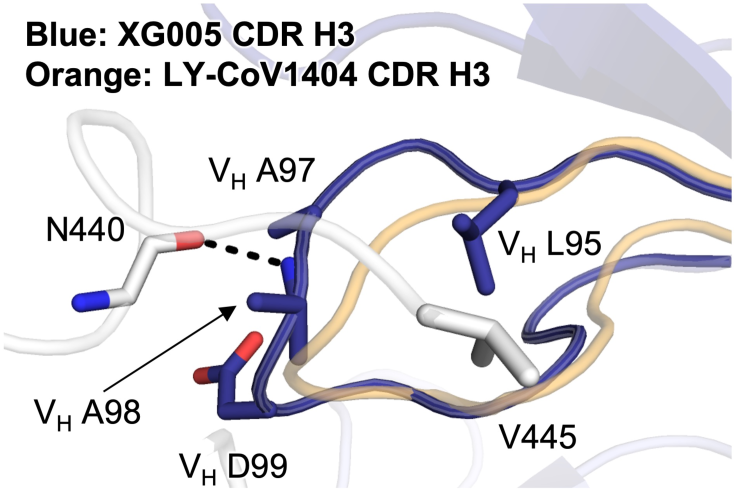
Interactions between RBD and CDR H3 of XG005. A cryo-EM structure of SARS-CoV-2 spike protein in complex with XG005 (PDB 7V26) that was reported in a previous study [20] is shown. A hydrogen bond between XG005 and the RBD is represented by a black dashed line. The CDR H3 of LY-CoV1404 (PDB 7MMO) [7] is also shown here as a transparent orange cartoon to demonstrate the relative positions of the CDR H3 loops from these two antibodies after superimposition.

